# New parameters for the in vitro development of cell lines from fish species

**DOI:** 10.1101/2023.05.09.539854

**Authors:** Ivanete de Oliveira Furo, Lygia S. Nogueira, Rodrigo Petry Corrêa de Sousa, Glaucia Caroline Silva-Oliveira, Diovanna Mirella dos Santos da Silva, Allan Costa-Malaquias, Edivaldo H.C de Oliveira

## Abstract

The establishment of cell lines from fish species is becoming important because of the increase in interest of these cells for viral analysis, environmental monitoring and cytogenetic studies. However, there is some discussion about the best conditions for maintaining these cells. We describe here a protocol for primary cell lines which we have found to be reliable. Fin biopsies from two species, *Geophagus proximus* and *Astyanax bimaculatus*, were isolated and cultured. We used three culture media (Leibovitz-L15, Dulbecco s Modified Eagle Medium-DMEM and 199) with or without the addition of AmnioMax and a standard temperature of 29ºC. The results showed that 199 medium was less efficient for both species. However, the cells of *G. proximus* had better proliferation in DMEM and L-15 media, while *A. bimaculatus* cells fared better in DMEM medium. The high concentration of L-glutamine and branched-chain amino acids (BCAAs) in the DMEM and L15 media was probably important for better adaptation of these cells. Furthermore, the addition of AmnioMax, a supplement rich in L-glutamine, increased cell proliferation in both species. Thus, the protocol initially established was tested in other tissues (fin, gills and kidney) of other fish species from the Amazon region. The cells were maintained in L-15 medium supplemented with 20% FBS (Fetal Bovine Serum) and 5% AmnioMax. It was observed that the cells were successfully subcultured and had a good proliferation, also the morphological characteristics were preserved. Thus, the methodology described in this research represents an innovative tool for the establishing of fish cell.

## 1. Introduction

Fish cells are excellent experimental models for a wide variety of studies. The first description of a fish cell line was made by Wolf & Quimby (1962), who established a cell line from *Salmo gairdneri* (Rainbow trout). Since then, other cell lines have been established and used as models in studies of embryology, toxicology, cytogenetics, virology, and environmental biology (Salvo et al., 2022; Fisher et al., 2003; Marques et al., 2007; Morcillo et al., 2016; Gjessing et al., 2018; Paim et al., 2018). The development of this technology has increased over recent years, due the greater use of animals in scientific investigations, especially for drug testing in the pharmaceutical industry (Doke & Dhawale, 2015).

Fish cell culture poses special difficulty, as these cells vary in the attachment intensity according to tissue and substrate used (Hightower & Renfro, 1988). Furthermore, the incubation temperature of fish cell culture can range from 4°C to 24ºC for species that live in cold water, or from 15ºC to 37ºC for species from tropical regions (Fryer & Lannan, 1994). Additionally, the requirements for culture media can vary according to the cell type. Complex media, such as L15, DMEM, 199, or RPMI 1460 are usually used together for the supplementation of different growth factors (Hightower & Renfro, 1988; Kuma et al., 2001; Lakra et al., 2011; Yashwanth et al., 2020). Nevertheless, only a few studies have compared the proliferative behaviour of fish cells in different culture media (Cheng et al., 1993; Kumar et al., 2001).

Given these aspects, this study presents a more comprehensive analysis of the optimum conditions for the in vitro culture of fish cells from the Amazon region. We evaluate the proliferative behaviour of cells from fins of *Geophagus proximus* (Castelnau, 1855) and *Astyanax bimaculatus* (Linnaeus, 1758). Additionally, the following parameters are considered: A) Can a standard protocol be established for fish cells line from the Amazon region? B) Does the addition of AmnioMax® in fish cell cultures improve cell growth?

## 2. Materials

### 2.1 Chemicals

- Dulbecco’s Modified Eagle Medium (DMEM), powder with high glucose (Cat. Nº: 12800017, Gibco, USA);
- Leibovitz’s L-15 Medium, powder (Cat. Nº 41300070, Gibco, USA);
- Medium 199, Earle’s Salt, powder (Cat. Nº31100035, Gibco, USA);
- Penicillin-Streptomycin Solution (Cat. Nº 15140130, Gibco, USA);
- Fetal Bovine Serum (Cat. Nº 26140079, Gibco, USA)
- Collagenase, Type IV, powder (Cat. Nº17104019, Gibco, USA);
- AmnioMAX™-II Complete Medium (Cat. Nº11269016, Gibco, USA);
- Trypan Blue Solution, 0,4% (Cat. Nº 15250061, Gibco, USA);
- EDTA-Trypsin (Cod 7720, Inlab, Brazil);
- Dimethyl sulfoxide (Cod 101862163, Sigma Life Science, France);
- 3-(4,5-Dimethylthiazol-2-yl)-2,5-Diphenyltetrazolium Bromide (MTT) (Cod 7318, Inlab, Brazil);
- Colchicine (Cat. Nº 77120, Serva Electrophoresis, Germany);
- Methanol (Cod 33209, Sigma-Adrich, France);
- Acetic acid glacial (Cod 33209, Sigma-Aldrich, France);
- Giemsa Stain (Cod 1.09203.0025, Merck, Germany);
- Ethanol (Cod 1.00983, Merck, Germany);

### 2.2 Equipment

- Incubation (Shel Lab, USA);
- Laminar flow Cabinet (ESCO, Brazil);
- Mr. Frosty (Cat. Nº 5100-0001, Nalgene, Germany);
- Ultrafrezeer -80 (Indrel, Brazil);
- Phase Contrast Microscope (Zeiss, Germany);
- Centrifuge (Novatecnica, Brazil);
- GloMax Multi detection System (Promega, USA);
- Microscope Leica DM1000 (Leica Microsystems, USA);
- GenASIs Software (DS Simage, ASI, USA);
- Software SigmaPlot 11.0 (Systat Software, SigmaPlot, UK);

## 3. Methods

### 3.1 Animal samples

For this study, we collected samples from two species of fish: *G. proximus* from Furo das Marinhas, situated in the Municipality of Benfica (Pará, Brazil, 1º17’28.3668” S, 48º19’41.1348” W). *A. bimaculatus* from Lagoa Salina, situated in the Caeté River Estuary (Pará, Brazil, 0º53’49.5” S, 46º40’05.9” W). The experiments followed ethical protocols and were approved by the ethics committee (CEUA-Federal University of Pará) under no. 9847301017/2018 and ICMBIO/SISBIO: 60197/2017.

### 3.2 Sample separation

Initially, using a sterile scalpel blade, the anterior fin of the two species mentioned above was collected. Then, the samples were washed in 1% bleach (NaOCL) and 70% alcohol. Finally, they were transported to the Laboratory of Cytogenomics and Environmental Mutagenesis, Evandro Chagas Institute (IEC), in DMEM medium, supplemented with 1% penicillin-streptomycin and 10% Fetal bovine serum (FBS).

### 3.3 Tissue decontamination and cell isolation

The collected tissue samples were decontaminated using the following steps: 8 seconds in Hank’s solution supplemented with 1% antibiotic; 2 seconds in 70% alcohol; 15 seconds in 1% bleach; 8 seconds in 70% alcohol. Then, finally the sample were processed in L-15 medium with 20% Fetal Bovine serum and 1% penicillin-Streptomycin.

Subsequently, the samples were cut with scalpels into smaller pieces, and incubated with collagenase type IV (0.0465 g/mL DMEN) for 45 minutes at 37ºC in 0,5% CO_2_. Throughout this period, the tissues were homogenised to accelerate the dissociation process. Afterwards, 5mL of L-15 medium were added to each tube followed of centrifugation for 10 minutes at 1000 rpm. Next, the supernatant was discarded, and the cell pellet was resuspended in fresh culture medium and centrifuged for 10 minutes at 1000 rpm. Finally, 5mL of L-15 medium supplemented with 20% FBS and 5% AmnioMax® were added. Cells were seeded on 25-cm^2^ tissue flasks and cultured at 29ºC without CO_2_.

Cell growth was monitored daily using a phase-contrast microscope (Zeiss, Germany) attached to a CCD camera (Leica DM1000, USA). After cell growth had reached 70–80% confluence, cells were detached by 1 ml of 0.025% Trypsin–EDTA (37ºC) and subsequently seeded into 75-cm^2^ tissue flasks in L-15 medium with 20%FBS. Cultures were continued up to the seventh passage.

### 3.4 Cryopreservation

When confluence was reached, the cells were removed with 0.025% Trypsin– EDTA, washed in Hank’s solution and centrifuged for 10 minutes at 1000 rpm. The isolated cells were suspended at 10^7^ cells/mL in cryopreservation solution (10% Dimethyl sulfoxide/ DMSO + 90% FBS). Next, aliquots of 1.5 mL of suspended cells were placed into 2 ml cryovials and kept at -80ºC for 24h. After this, the cryovials were transferred to liquid nitrogen for long-term storage. To test whether cryopreservation affected the phenotype and growth, some cryovials of both species were thawed by the fast method. Briefly, vials were removed from liquid nitrogen and placed in a 37°C water bath for one minute. Next, cells were seeded into 75-cm^2^ tissue flasks in L-15 medium with 20%FBS and 5% AmnioMax®.

### 3.5 Experimental conditions

For this study, L-15, DMEM, and 199 media with/without AmnioMax were used to compare the outcome of cell culture in G. *proximus* and *A. bimaculatus* (Table 1) The concentration of AmnioMax® was based on Furo et al. (2017).

**Table 1.** Conditions used for the fin cells of G. proximus and A. bimaculatus.

| Type of treatment | Media | Growth facts | Temperature | CO<sub>2</sub> |
| --- | --- | --- | --- | --- |
| **A** | L-15 | 20% FBS | 29°C | Absence |
|  | DMEN |  |  |  |
|  | 199 |  |  |  |
| **B** | L-15 | 20% FBS + 5%<br>AmnioMax® | 29°C | Absence |
|  | DMEN |  |  |  |
|  | 199 |  |  |  |

#### 3.5.1 Cell Viability Assay

The trypan blue exclusion assay was used to quantify cell viability. In this process, 5 × 10^4^ cells were seeded into 6-well plates, and the percentage of live cells determined after 24h and 72h of culture (using condition A and B, as described in Table 1). Cells were counted by light microscopy using a Neubauer hemocytometer. The experiments were performed in triplicate and repeated at least three times.

#### 3.5.2. Evaluation of Mitochondrial Activity

The mitochondrial activity of viable cells was evaluated by the compost reduction 3-[4,5-dimethylthiazol-2-yl]-2,5-diphenyl tetrazolium bromide (MTT) procedure (Mosmann et al., 1983). The cells were plated in 24-well plates at a density of 1×10^4^ cells/well in 100µL of medium (using conditions A or B, as described in Table 1). After 24h and 72h, the cells were washed once with Hank’s solution, and the MTT assay was performed following the manufacturer’s instructions. The intensity of formazan staining was measured at 560 nm by the GloMax Multi detection System (Promega). The results are represented as relative light units (RLU) for viable cells (RLU/ viable cells).

#### 3.5.3 Analysis of ATP Production

The ATP production was assessed by the Mitochondrial ToxGlo™ Assay kit following the manufacturer’s instructions (Promega, Germany). Briefly, cells were plated (1.5×10^3^ cells/well) (using conditions A as described in Table 1) and further lysed. After 72h treatment, Cell lysate was diluted 1:10 and incubated with ATP Detection Reagent. The absorbance was read at 595 nm using the GloMax Multi detection System (Promega, USA).

### 3.6 Evaluation of proliferative potential and cell morphology

All cells were cultured separately in 25cm^2^ tissue culture flasks using different culture media (DMEM, L15 or 1999) supplemented with 20% FBS at 29°C. The cells were monitored daily using a phase-contrast microscope (Zeiss, Germany) with attached CCD camera (Leica DM1000, USA). The cell proliferation was monitored daily and alterations in morphological pattern were recorded.

### 3.7 Chromosome preparation

Mitotic chromosomes were obtained by adding 0.016% colchicine (100 µL for each 5 mL of culture medium) in culture flasks that showed an approximate 50% mitotic index. These flasks were incubated for 4 hours at 29ºC without CO_2_ buffering. The cells were removed using 0.025% EDTA-Trypsin and centrifuged at 2000rpm for 10 minutes, followed by hypotonic treatment (0.075M KCl) at 37ºC for 1h 20 before fixation in Methanol: Acetic acid (3:1 v/v).

At least 15 metaphases in conventional staining (Giemsa 5% in phosphate buffer, pH 6.8) were examined to determine the diploid number and karyotype of each animal investigated. Images were captured using a 100× objective (Leica DM1000) and GenASIs software (ADS Biotec).

### 3.8 Statistic Analysis

Cell numbers were determined after three passages, and median and standard deviations obtained. Comparison between the treatments was carried out by two-way analysis of variance (ANOVA), followed by the Tukey test. In all cases, the significance level adopted was 95% (α = 0:05). Statistical analyses were performed using SigmaPlot 11.0 (Systat Software, SigmaPlot, UK).

### 3.9 Application of the established protocol

The cell growth condition initially established (L-15 media, 20% FBS and 5% AmnioMax^®^) were tested in different types of tissues of the fish species from Amazon region (Table 2). Cells were seeded on 25-cm^2^ tissue flasks and cultured at 29ºC, without CO_2._ The cells growth was monitored daily using a phase-contrast microscope (Zeiss, Germany) with attached CCD camera (Leica DM1000, USA).

**Table 2.** Species and Tissues used in the experiments

| **Species Name** | **Type of tissue** | **Collect place** |
| --- | --- | --- |
| *Guavina guavina* | fin and kidney | Lagoa Salina, in the Caeté River Estuary/PA |
| *Bagre bagre* | gills | Lagoa Salina, in the Caeté River Estuary/PA |
| *Conodon nobilis* | fin | Lagoa Salina, in the Caeté River Estuary/PA |
| *Astyanax scabripinis* | fin | Lagoa Salina, in the Caeté River Estuary/PA |

## 4. Results

### 4.1 Development of cell line

Fin cells from *A. bimaculatus* and *G. proximus* attached to the culture flask surface within 4 hrs post incubation. The cells reached 85–90% confluence within the seventh day, and then passaged into new culture flasks with fresh medium. The cells were dissociated within 1 min of trypsin addition. Cell replication was slow at the initial stages of culture, but this improved with later passages.

### 4.2 The Morphology and Density of cells

The analysis of cultures by phase contrast microscopy showed that the morphological characteristics of cells of both species (*A. bimaculatus* and *G. proximus*) were preserved in all media tested at 24h and 72h. The cells have elongated and fusiform features like other fibroblast cells described in the literature (Fig. 2).

**Figure 1.**
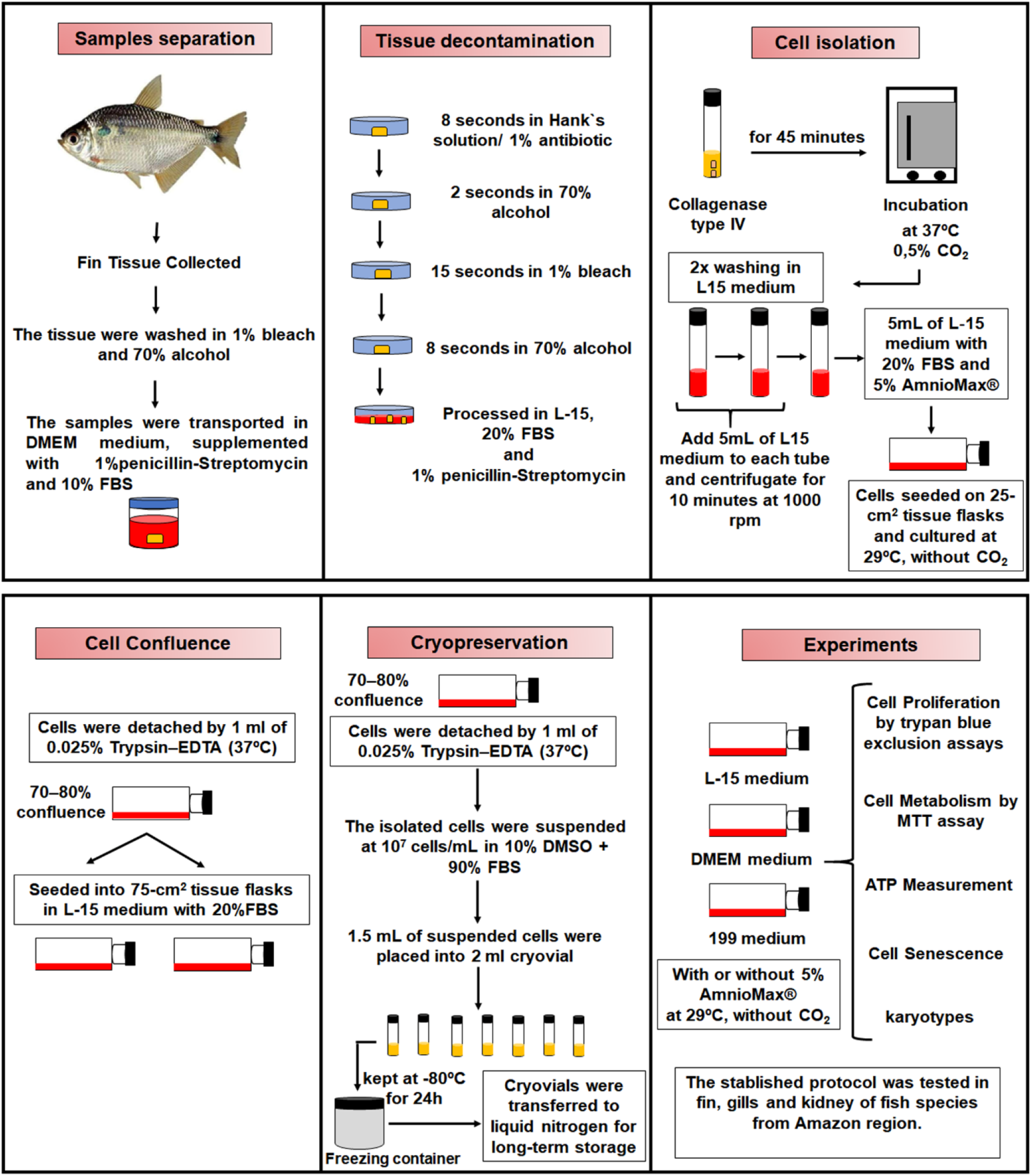
Protocol followed to initiate of cell culture

**Figure 2.**
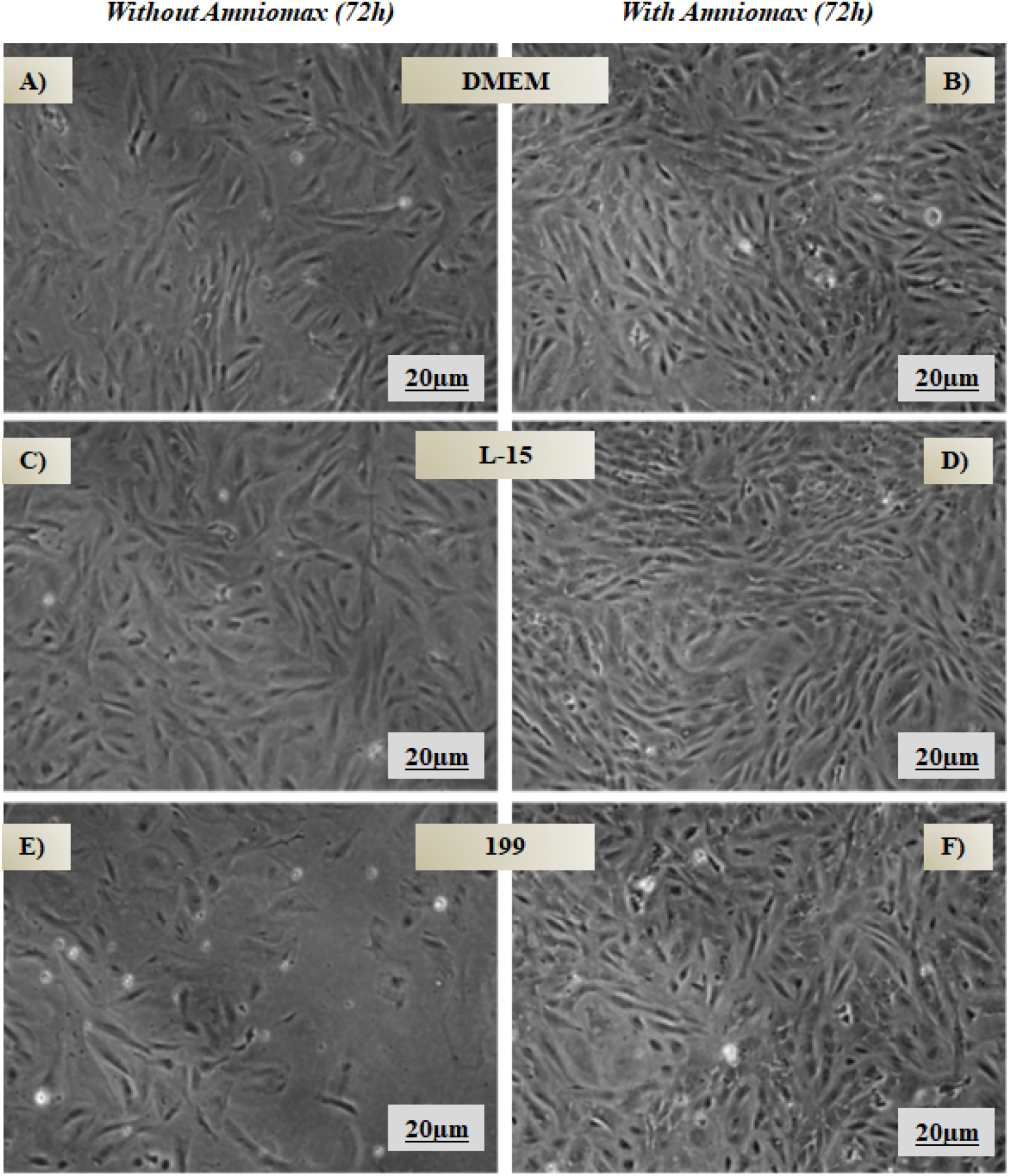
Fin cells of *A. bimaculatus* in 24 and 72h. Without AmnioMax^®^ (A, C and E). With AmnioMax^®^ (B, D and F). Bar=100µm

### 4.3 Cell thawing

The thawed cells showed viability even after months in liquid nitrogen. The average estimated revival percentage was 60% of the initial cell population. The revived cells recovered well and grew to confluency within 7 days.

### 4.4 Cell Proliferation by Trypan Blue Exclusion Assay

The number of viable cells in culture of *A. bimaculatus* cell was similar in all media after 24h. However, a significant reduction of viable cells was observed after 72h in the culture with 199 medium (Fig. 3A). Moreover, in *A. bimaculatus* cells, the addition of 5% AmnioMax^®^ in all media resulted in similar cell proliferation after 24h. Nevertheless, in the treatments with AmnioMax^®^ for 72h, the cells cultured in DMEM and 199 media significantly increased in number (Fig. 3C).

**Figure 3.**
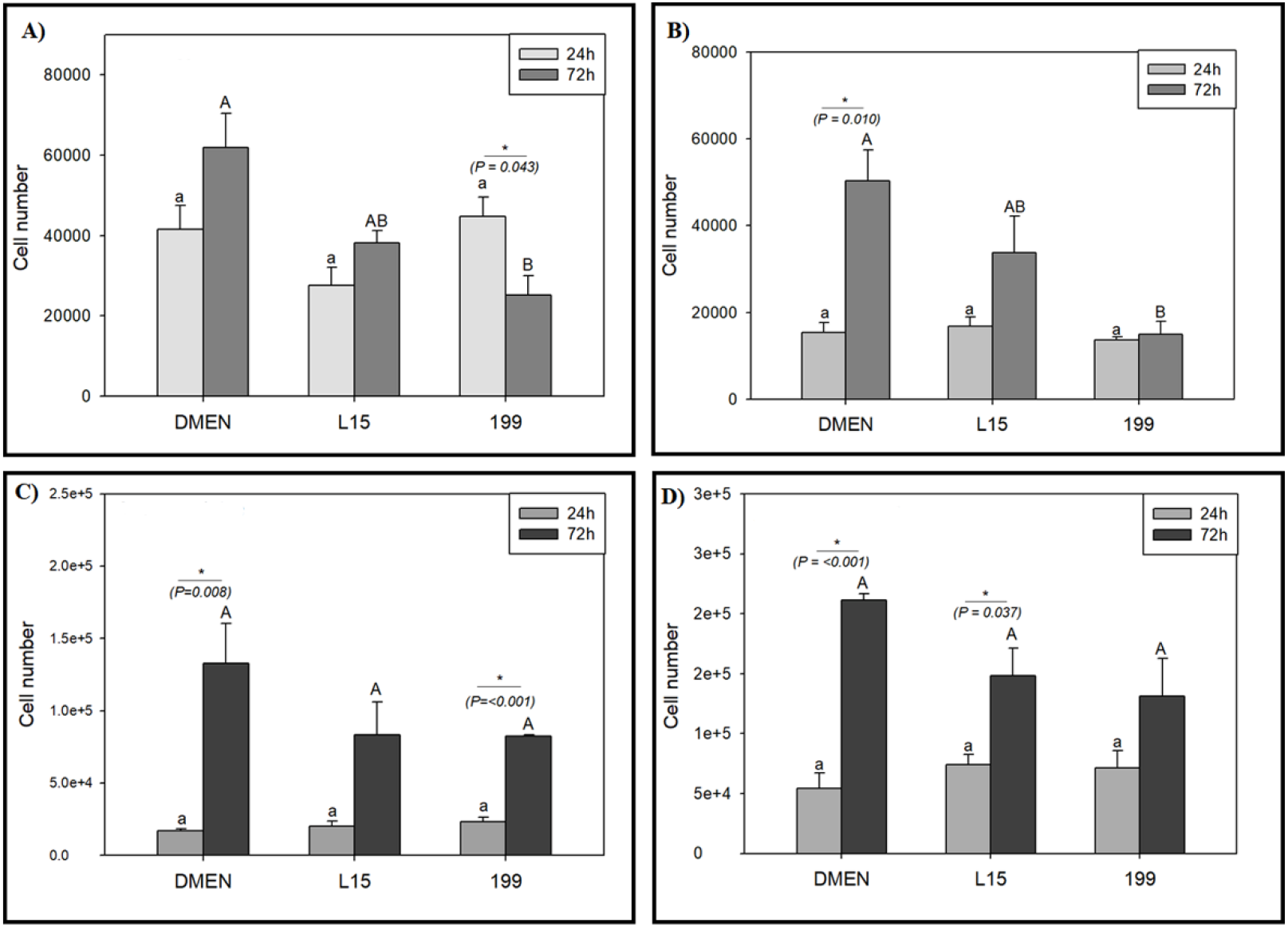
Cell proliferation of *A. bimaculatus* and *G. proximus*. Without AmnioMax^®^ (A, B). With 5% AmnioMax^®^ (C, D). **Legends:** Lowercase, represents statistical difference between the treatments at 24h and uppercase difference between the treatments at 72h. (*) difference between 24h and 72h in the same culture medium.

Furthermore, *G. proximus* cells maintained similar their proliferative for 24h regardless of the type of media used (Figure 3B). However, for 72h we observed that cells cultured in DMEM medium significantly increased their proliferation potential. Interestingly, for 72h the addition of 5% AmnioMax® the cultures of *G. proximus*, significantly increased their proliferation in the DMEM and L15 media (Fig. 3D).

### 4.5 Mitochondrial Activity

The highest cellular metabolism of *A. bimaculatus* was observed in L15 medium, followed by DMEN and 199 media at 24h of culture (Figure 4A). On the other hand, the metabolism rates at 72h reached the highest values in the 199 medium. The presence of 5% AmnioMax^®^ in the culture of *A. bimaculatus*, maintained similar cell metabolism rates in all media at 24 and 72h (Fig. 4C).

**Figure 4.**
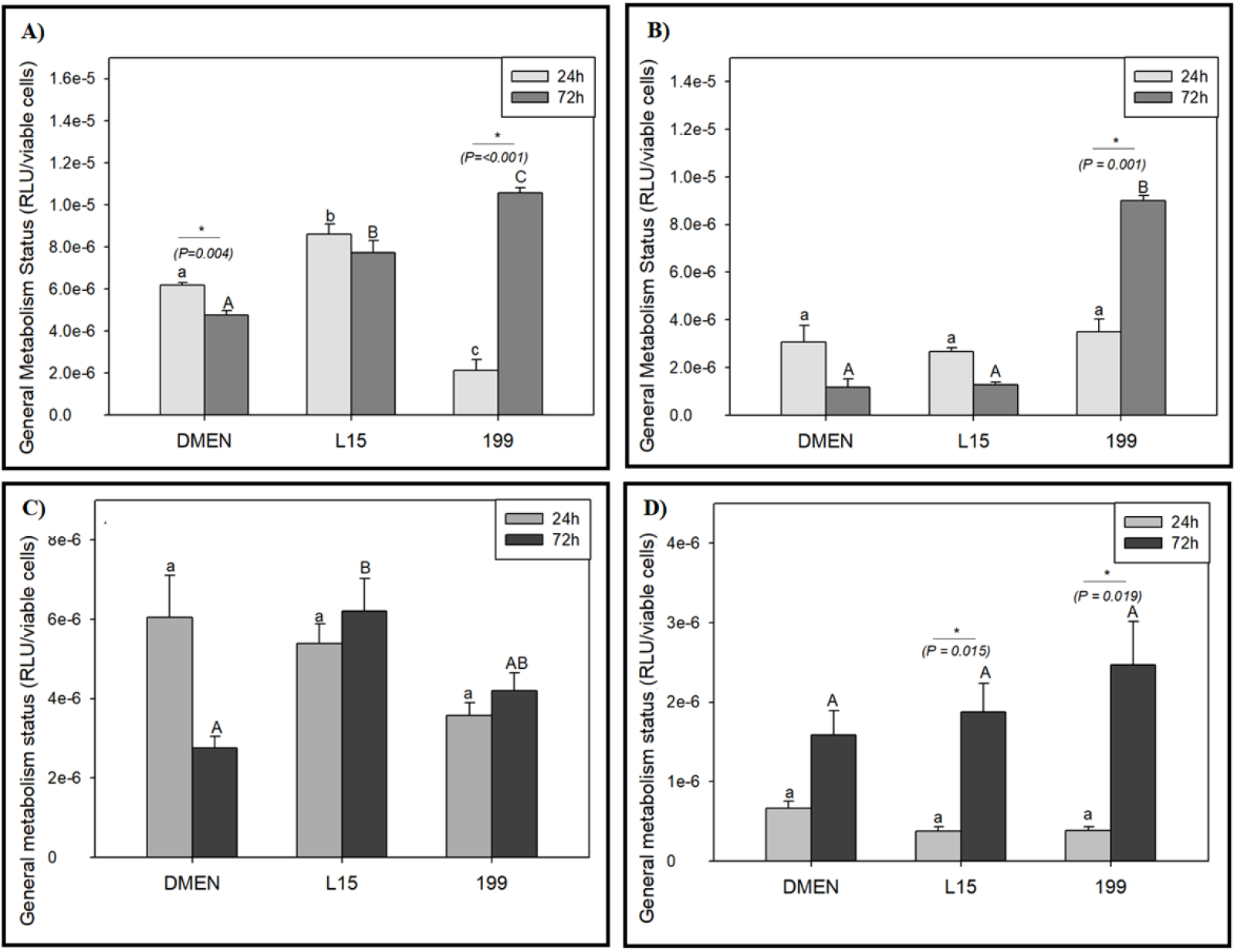
Cell metabolism of *A. bimaculatus* and *G. proximus*. Without AmnioMax^®^ (A, B). With AmnioMax^®^ (C, D). **Legends:** Lowercase represents statistic difference between the treatments at 24h. Uppercase represent difference between the treatments at 72h. (*) Statistical difference between 24h and 72h in the same culture media.

On the other hand, *G. proximus* cells exhibited similar metabolism in all culture media at 24h. However, at 72h a significant increase in cell metabolism was observed in 199 media (Figure 4B). The addition of 5% AmnioMax^®^ in the *G. proximus* cells, resulted in a significantly increase of cell metabolism at 72 h in the L-15 and199 media (Fig. 4D).

### 4.6 ATP Production

The results of mitochondrial activity in 72h without the presence of AmnioMax^®^ in both fish (*A. bimaculatus* and G. *proximus*) support the results of general metabolic activity. Cells from *A. bimaculatus* show decreased ATP production in the DMEM medium compared to the L-15 and 199 media. On the other hand, the cells from *G. proximus* achieved the highest ATP production only in the 199 medium (Figure 5).

**Figure 5.**
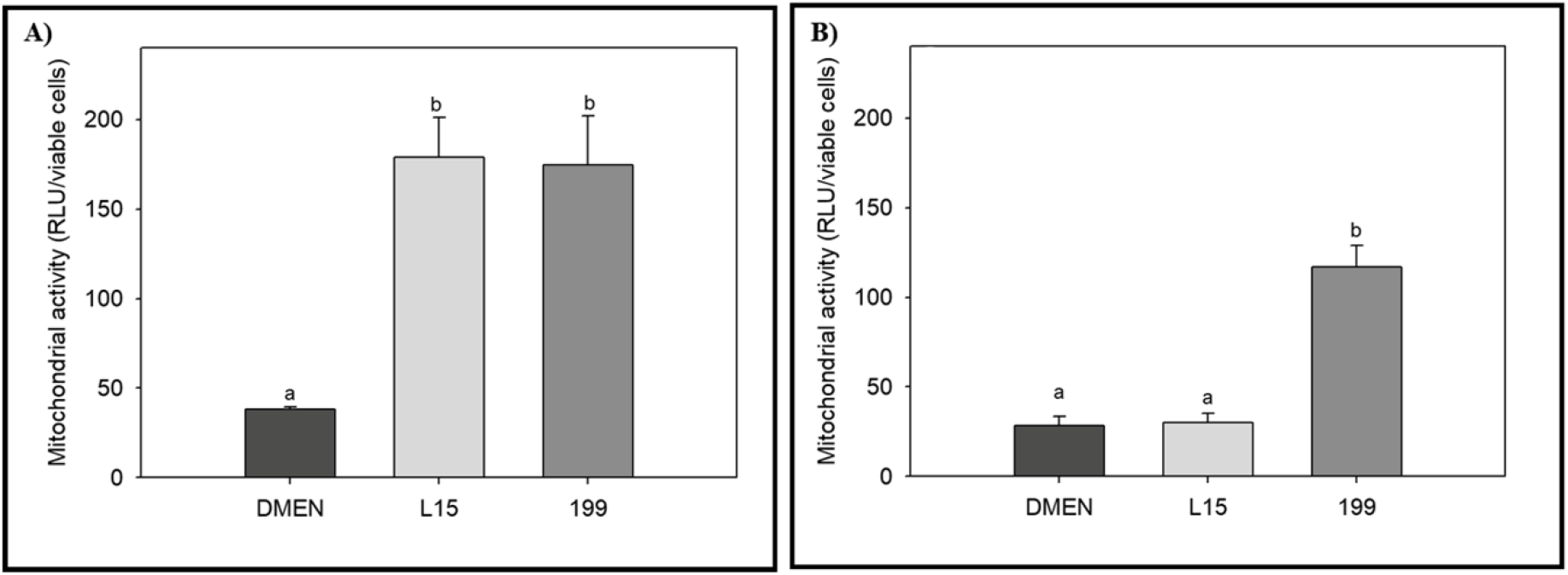
ATP production in primary culture of *A. bimaculatus* (A) and *G. proximus* (B). **Legend:** Lowercase represents statistical difference between the culture media.

### 4.7 Alterations in Survival

The *A. bimaculatus* cells cultured with DMEM showed stable growth until the nineteenth passage. After that, the cells grew rapidly, which prompted us to maintain them with only 10% FBS. These cells overpassed the #60 passage and were frozen and stored in liquid nitrogen. *A. bimaculatus* cells cultured in L-15 and 199 media survived until the #26 and #7 passage, respectively. The DMEM-cultured cells of *G. proximus*, died at #18. However, on L-15 and 199 media, they became senescent at #14 and #6, respectively (Table 3).

**Table 3.** Cell senescence. The passage number are represented by (#)

| Species name | Media/ Passages of cell senescence |  |  |
| --- | --- | --- | --- |
|  | DMEN | L15 | M199 |
| *A. bimaculatus* | #60> | #26 | #7 |
| *G. proximus* | #18 | #14 | #6 |

### 4.8 Chromosome Analysis

The chromosome counts of 15 metaphase plates revealed a diploid number at the 4 and 6th passage of *A. bimaculatus* and *G. proximus* cells of 50 and 48, respectively. Their karyotypes are shown in Fig. 6.

**Figure 6.**
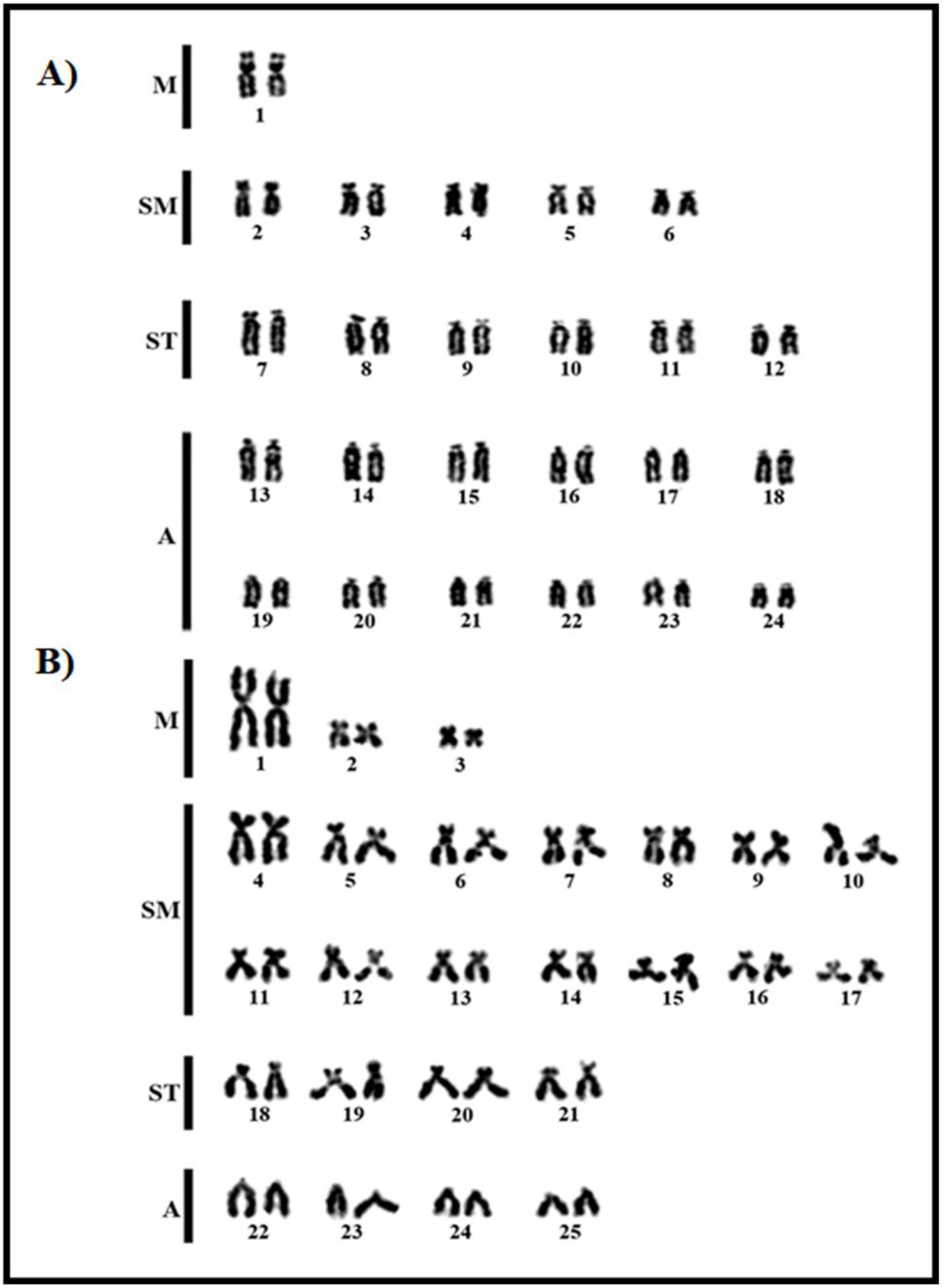
Karyotype with conventional staining (A) *G. proximus* (2n=48) and (B) *A. bimaculatus* (2n=50).

### 4.9 Application of the established protocol to other species

The cells of fin, kidney and gills isolated from other fish species from the Amazon region grew well and had well preserved morphology; the cells were elongated and fusiform (Fig.7), similar to other fibroblasts. They reached 80% confluence within 7 days (Fig.7).

**Figure 7.**
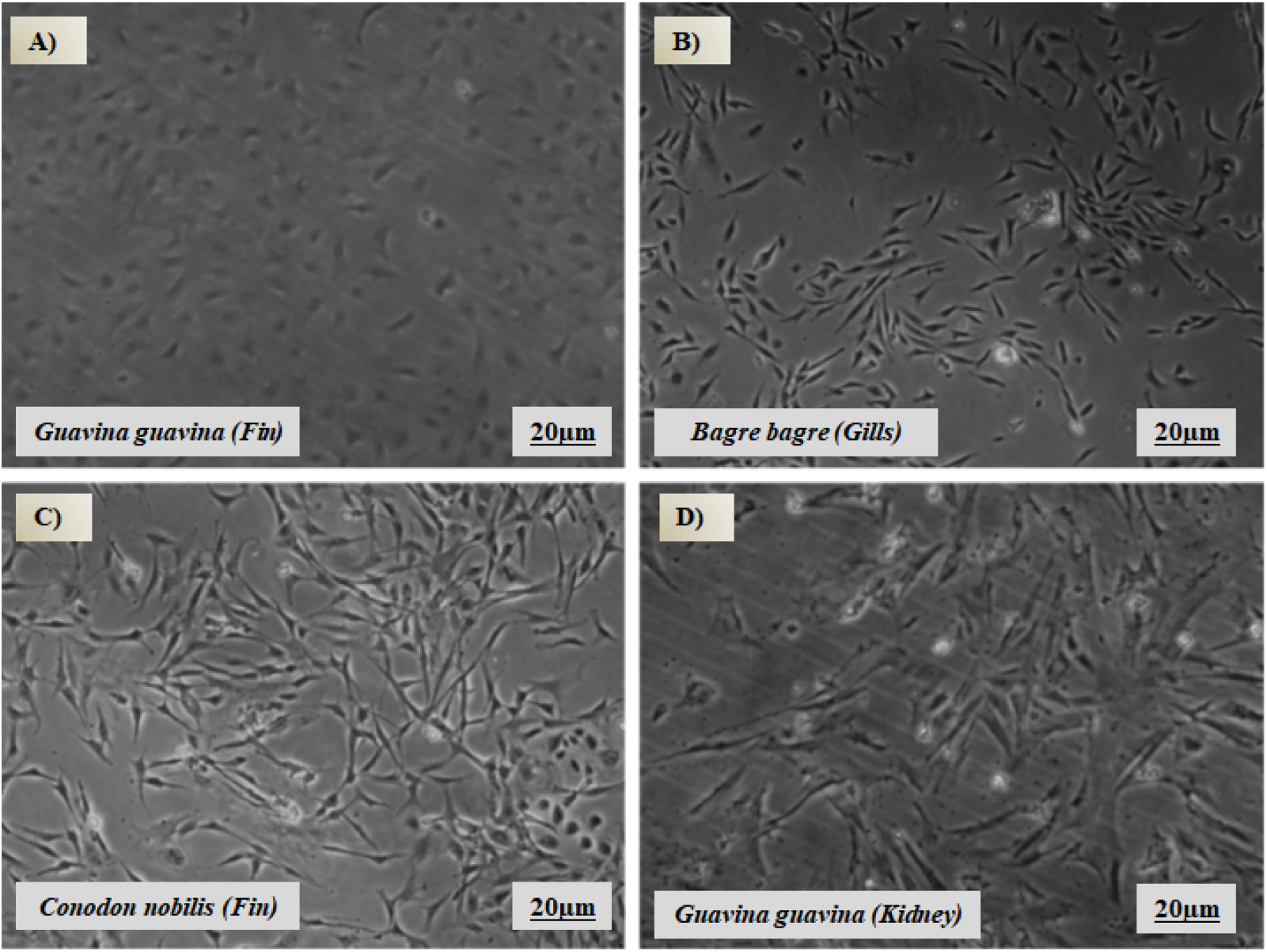
Fish cell proliferation by phase contrast microscope. Bar=20µm

## 5. Discussion

### 5.1 Physiological Parameters

An incontestable point in cell culture of fish species is the choice of culture media (Yao & Asayama, 2017). Our results show that the 199 medium was less efficient in both species. At 72h, there is a decrease and/or an absence of proliferating cells and a marked increase in metabolic activity (Fig. 3A-B, 4A-B and 5). The increased metabolism is related to an increase in total energy expenditure to maintain vital activity. According to Diaz and Vazquez (2018), the required energy increases beyond the basal metabolic rate when the cell grows or performs other functions. Thus, despite the 199 medium used in some fish cell lines (Cheng et al., 1993), our evaluation shows that this medium is inadequate for the *G. proximus* and *A. bimaculatus* cells.

Furthermore, for the L-15 and DMEM media, *G. proximus* cells show a lower metabolic rate and good proliferative potential when compared to *A. bimaculatus* cells, in which the metabolism rate was significatively reduced only in DMEM media (Fig. 3A-B, 4A-B). Additionally, analysis of ATP production confirms the results of general metabolism (Fig. 5). The results indicate that *G. proximus* cells proliferate favorably in L-15 and DMEM media, whereas this is observed only in *A. bimaculatus* cells in DMEM medium.

The fish cultures reported in other studies present a great diversity in culture media. However, the cells were mainly cultured as DMEM or L-15 media (Lakra et al., 2011). Despite that, some cells, as in *Astyanax* sp. (Paim et al., 2018), adapt only to DMEM medium with its complex chemical composition, and high concentrations of amino acids, vitamins, and glucose (Yao & Asayama, 2017).

A comparation of the chemical composition of culture media in *Astyanax* sp. (Paim et al., 2018), show a higher concentration of L-glutamine in DMEM and L-15 than in 199 medium. The non-essential amino acids play different roles. (e.g., metabolism, cell proliferation, protein synthesis). Cell stress can lead to the lower availability of these amino acids, leading to decreased resistance of cells to injury, thus inducing apoptosis (Ko et al., 2001; Curi et al., 2007). It is possible that the high concentration of L-glutamine in DMEM and L-15 media was probably a factor important for the proliferation of *G. proximus* and *A. bimaculatus* cells. The lower availability of this amino acid could still explain the absence of proliferation and the death of cells maintained in 199 medium.

Another possibility is the variation in high concentrations of other amino acids in DMEM and L-15 media, such as the branched-chain amino acids (BCAAs) valine, leucine, and isoleucine, which are widely studied due to their effect in metabolism, cell growth and oxidative stress responses (Shao et al.,2018; Sartori et al., 2020). Maybe, the combination and high concentration of these amino acids (L-glutamine and branched-chain amino acids) in L-15 and DMEM media are important in the culture of *G. proximus* and *A. bimaculatus* cells.

### 5.2 AmnioMax^®^ as a growth factor

Fish cells require supplementation with growth factors other than Fetal Bovine serum, as demonstrated by Kuma et al., (2001). In this study, the authors established cultures from ovary tissues of African catfish using many additives, including fish muscle extract, sucrose, and prawn shell extract. Epidermal growth factor (mEGF) (Watanabe et al., 1987) and basic fibroblast growth factor (bFGF) (Chen et al., 2004) have also been used as additives.

The AmnioMax^®^ was initially formulated for primary cultures of amniotic cell fluid used in prenatal diagnosis. Fetal Bovine serum and L-glutamine are included in AmnioMax^®^ to maximize cell attachment and proliferation (www.lifetechnologies.com/support). Our studies report the use of AmnioMax^®^ as an additive for fibroblast culture growth in birds (Furo et al., 2017; 2020). However, studies involving AmnioMax^®^ in the proliferation of fibroblast culture from fish are unknown until the present. Our analyses demonstrate that AmnioMax^®^ supplementation increases cell proliferation in both species (Fig.3C and D). In *A. bimaculatus* a minimal effect of AmnioMax on cultures was observed, for the 199 and L-15 media (Fig. 3C).

In *G. proximus* cells, AmnioMax^®^ supplementation significantly increased cell metabolism in L-15 and 199 media (Fig. 4D). The increase could be explained by improved establishment in these cultures (Fig.3D). The *G. proximus* cells in L-15 medium proliferated intensely with AmnioMax^®^ compared to its absence (Fig. 3B e D). One aspect to be highlighted is that AmnioMax^®^ maximized cell attachment in all *G. proximus* cultures (Fig. 3B and D). However, this was not observed in *A. bimaculatus* cultures (Fig. 3A e C).

### 5.3 Cytogenetics Parameters

Metaphase chromosome spreads were made for karyotyping the cell lines using conventional staining.

The *G. proximus* and *A. bimaculatus* cells at the 6 and 4^th^ subculture, presented 2n=48 and 50, respectively, conserving the original karyotype of the species, both in diploid number and chromosomal morphology (Fig. 6) (Feldberg et al., 2003; Nishiyama et al., 2016). This confirms that these fish cell lines are reliable models for toxicology essays and studies of environmental biology (Sousa et al., 2022; Morcillo et al., 2016), important aspects for the reduction in the use of live animals in scientific investigations.

## 6. Conclusion

A standard protocol for maintaining most fish cell cultures is established. We suggest the L15 medium, supplemented with 20% FBS and 5% AmnioMax. As observed in Figure 7, the cells in L15 show excellent attachment and proliferation. However, for some species, such as *Astyanax* sp., we found better growth in DMEM medium (Paim et al., 2018), suggesting that where L15 is associated with poor growth, DMEM medium should be substituted.

The results obtained demonstrate that the concentration of L-glutamine and branched-chain amino acids in the chemical composition of culture media is probably an important factor in the establishment and cell proliferation of *G. proximus* and *A. bimaculatus*. Analyses with AmnioMax^®^ showed that the presence of this growth factor in culture potentiated cell proliferation and, in the culture of *G. proximus*, maximized cell attachment to the substrate. Thus, it is suggested that the innovative protocol described here can be applied successfully to fish studies, particularly to fish from the Amazon region.

## Contributions

Conceptualization, I.O.F. and E.H.C.O.; Data curation and formal analysis, I.O.F., L.S.N., A.C.M and E.H.C.O.; Investigation, I.O.F., L.S.M., L.S.N.; Methodology, I.O.F., L.S.N., S.R.C., D.M.S.S., G.C.S.O.; R.C.S Project administration, E.H.C.O.; Funding acquisition, E.H.C.O.; Validation, I.O.F. and E.H.C.O.; Writing (original draft), I.O.F. L.S.N and E.H.C.O.; Writing (review and editing), L.S.N., E.H.C.O.

## Acknowledgments

Authors would like to thank the staff of Laboratory of Cytogenomics and Environmental Mutagenesis, Evandro Chagas Institute (IEC), CAPES for technical and financial support and the Professor Malcolm A. Ferguson-Smith for review and English edition.

## References

1. Chen SL, Ren GC, Sha ZX. et al. Establishment of a continuous embryonic cell line from Japanese flounder Pararlichthys olivaceus for virus isolation. Dis Aquat Organ 60:241–246, 2004.

2. Cheng LL, Bowser PR, Spitsbergen JM. Development of cell cultures derived from lake trout fiver and kidney in a hormone supplemented, serum-reduced medium. Journal of Aquatic Animal Health. 5:119–126, 1993.

3. Curi R, Newsholme P, Procopio J. et al. Glutamine, gene expression, and cell function. Frontiers in Bioscience 12: 344–357, 2007.

4. Doke SK, Dhawale SC. Alternatives to animal testing: A review. Saudi Pharmaceutical Journal, 23: 223–229, 2015.

5. Feldberg E, Porto JIR, Bertollo LAC. Chromosomal changes and adaptation of cichlidae fishes during evolution. In: Val AL, Kapoor BG. (Ed.). Fish adaptation. Enfield: Science Publishers, Inc,. p. 285–308, 2003.

6. Fernandez-de-Cossio-Diaz J and Vazquez A. A physical model of cell metabolism. Scientific Reports, 8:8349, 2018.

7. Fisher S, Jagadeeswaran P, Halpern ME. Radiographic analysis of zebrafish skeletal defects. Dev Biol 264:64–76, 2003.

8. Fryer JL, Lannan CN. Three decades of fish cell culture: A current listing of cell lines derived from fishes. Journal of tissue Culture methods. 16:87 – 94, 1994.

9. Furo IO, Kretschmer R, dos Santos MS. et al. Chromosomal mapping of repetitive DNAs in Myiopsitta monachus and Amazona aestiva (Psittaciformes, Psittacid ae: Psittaciformes), with emphasis on the sex chromosomes. Cytogenet. Genome Res. 151:151–160. 2017.

10. Furo IO, Kretschmer R, O’Brien PC. et al. Chromosomal Evolution in the Phylogeentic Context: A Remarkable Karyotype Reorganization in Neotropical Parrot Myiopsitta monachus (Psittacidae). Frontiers in Genetics 11:721, 2020.

11. Gjessing MC, Aamelfot M, Batts WN. et al. Development and characterization of two cell lines from gills of Atlantic salmon. PLoS ONE 13(2): e0191792, 2018.

12. Ko YG, Kim EY, Kim T. et al. Glutamine-dependent antiapoptotic interaction of human glutaminyl-t RNA synthetase with apoptosis signal-regulating kinase 1. J Biol Chem 276: 6030–6036, 2001.

13. Kuma GS, Singh ISB, Philip R. Development of a cell culture system from the ovarian tissue of African catfish Clarias gariepinus. Aquaculture 194: 51–62, 2001.

14. Laka WS, Swaminathan TR, Joy KP. Development, characterization, conservation and storage of fish cell lines: a review. Fish Physiol Biochem. 37:1–20, 2011.

15. Marques CL, Rafael MS, Cancela ML. et al. Establishment of primary cell cultures from fish calcified tissues. Cytotechnology 55:9–13, 2007.

16. Morcillo P, Chaves-Pozo E, Meseguer J. et al. Establishment of a new teleost brain cell line (DLB-1) from the European sea bass and its use to study metal toxicology. Toxicology in vitro, 2016. In press.

17. Mosmann T. Rapid Colorimetric Assay for Cellular Growth and Survival: Application to Proliferation and Cytotoxicity Assays. Journal of lmmunological Methods 65: 55–63, 1983.

18. Nishiyama PB, Vieira MMR, Portos FE. et al. Karyotype diversity among three species of the genus Astyanax (Characiformes: Characidae). Braz.J. Biol, 76:360–366, 2016.

19. Paim FG, Maia L, Alvarenga FCL. et al. New Protocol for Cell Culture to Obtain Mitotic Chromosomes in Fishes. Methods and protocols 47:1–9, 2018.

20. Hightower LE and Renfro LJ. Recent Applications of fish Cell culture to Biomedical research. The Journal of Experimental Zoology. Connecticut. 248: 290–302, 1988.

21. Sartori T, Santos ACA, Oliveira R. et al. Branched chain amino acids improve mesenchymal stem cell proliferation, reducing NF_B expression and modulating some inflammatory properties. Nutrition, 2020.

22. Shao D, Villet O, Zhang Z. et al. Glucose promotes cell growth by suppressing branched-chain amino acid degradation. Nature Communications. 9:2935, 2018.

23. Sousa AH, Pereira JPG, Malaquias AC. et al. Intracellular accumulation and DNA damage caused by methylmercury in glial cells. J Biochem Mol Toxicol. e23170:2022.

24. Watanabe T, Nakano M, Asakawa H. et al. Cell culture of rainbow trout liver. BJpn Soc Sci Fish 53:537–542, 1987.

25. Wolf K and Quimby MC. Established eurythermic line of fish cells in vitro. Science 135:1065–1066, 1962.

26. Yao T and Asayama Y. Animal-cell culture media: History, characteristics, and current issues. Reprod Med Biol.16:99–117, 2017.

27. Yashwanth BS, Goswami M, Valapiil RK. et al. Characterization of a new cell line from ornamental fish Amphiprion ocellaris (Cuvier, 1830) and its susceptibility to nervous necrosis virus. Scientific Report. 10:20051, 2020.

